# DAPL1 deficiency impairs autophagy in retinal pigment epithelium to drive age-dependent retinal pathologies

**DOI:** 10.1101/2025.06.09.658638

**Authors:** Huaicheng Chen, Qiufan Tan, Yishan Hu, Liping Liu, Xiaoxin Liang, Ling Hou

**Author notes:** These authors contributed equally to this work (Huaicheng Chen, Qiufan Tan).

## Abstract

Age-related retinopathy, primarily comprising age-related macular degeneration (AMD), is a major cause of irreversible blindness in the elderly population worldwide. Although its precise mechanisms remain incompletely understood, autophagy deficiency in retinal pigment epithelium (RPE) cells has been associated with AMD in patients; however, the regulatory pathways involved are not fully elucidated. In this study, we report that deficiency in the *Dapl1* gene inhibits RPE cell autophagy and leads to age-dependent retinal pathologies in mice.

Specifically, 18-month-old *Dapl1-/-* mice manifest age-related retinal dysfunction and structural abnormities, including increased retinal stress, photoreceptor/RPE cell damage, lipid aggregates and microglial activation. Furthermore, we demonstrate that autophagy is impaired in the RPE cells of *Dapl1-/-* mice, while overexpression of DAPL1 in RPE cells enhances autophagy activity. Mechanistically, we elucidate that DAPL1 suppresses the expression of E2F1 and c-MYC, which consequently downregulates mTOR and upregulates ATG16 and Beclin1 expression via DAPK1 in RPE cells, thereby promoting autophagy. Taken together, our findings demonstrate that DAPL1 acts as a novel regulator of autophagy in RPE cells, and its deficiency increases susceptibility to age-dependent retinal pathologies in mice.

## Introduction

Older adults exhibit heightened susceptibility to various ocular pathologies [1]. Among these, age-related macular degeneration (AMD) is the leading cause of irreversible blindness, which is closely related with the dysfunction of RPE cells [2–3]. This devastating disease affects millions worldwide, with approximately 80% of clinical cases presenting as dry AMD (geographic atrophy), characterized by progressive degeneration of RPE cells and subsequent photoreceptor damage [4]. As multifunctional retinal support cells, RPEs perform critical physiological roles including: (1) oxidative stress regulation through reactive oxygen species (ROS) clearance, (2) paracrine secretion of essential growth factors, (3) maintenance of blood-retinal barrier integrity, and (4) phagocytic removal of metabolic waste [5–6]. Both basic and clinical studies implicate the RPE cells act as a primary site in the retinopathies, while mutations in the RPE functional genes were demonstrated to induce retinal degeneration and visual lost [7–9]. Notably, age-dependent RPE senescence and dysfunction were believed to be the main causing factors of age-dependent retinal pathologies [10–11]. However, the precise molecular mechanisms underlying RPE senescence and its causal relationship with age-dependent retinal pathologies remain incompletely understood, warranting further investigation into the cellular aging processes and associated signaling pathways.

The pathogenesis of age-dependent retinal pathologies is mechanistically linked to chronic oxidative stress. Elevated levels of reactive oxygen species (ROS) induce macromolecular damage through peroxidation of lipid membranes, protein carbonylation, and nucleic acid fragmentation, triggering pathogenic aggregation of oxidized biomolecules. The age-progressive accumulation of these oxidatively modified macromolecules constitutes a key molecular pathway driving the pathogenesis of age-related retinopathies [1,12]. How to efficiently remove cellular waste in retinal cells is critical for keeping retinal homeostasis and preventing disease progression. In RPE cells, proteasomal degradation, heterophagy and autophagy play critical roles in clearing cellular waste. Among them, autophagy is a biological process to degrade the misfolding or unneeded proteins and damaged organelle in the lysosomes, which is necessary to protect the cells from stress induced injury and support energy [13–14]. Autophagy activity is regulated by the mTOR (mechanistic target of rapamycin) pathway [15], and involves numerous proteins, such as microtubule-associated protein light chain 3 (LC3), autophagy related proteins (ATGs) and Beclin1 [16].

It was reported that autophagy was impaired in RPE cells from AMD patients compared to healthy controls [17–18]. In mouse models, knockout of *Atg5*, *Atg7* and *Rblcc1*—disrupting autophagy—induces AMD-like phenotypes, including retinal generation, RPE cell damage and abnormal microglial accumulation [19–20]. Similarly, A2E-mediated inhibition of autophagy causes RPE cell damage [21]. Furthermore, autophagy is implicated in oxidative stress and inflammation [22], key initiating factors in AMD. Critically, the phagocytosis of shed photoreceptor outer segments (POS) by RPE cells—essential for retinal health—requires subsequent autophagy-lysosomal degradation [23–24]. These findings collectively indicate that autophagy is critical for retinal homeostasis and function. While Deficiency in RPE cell autophagy is linked to age-related retinal degeneration, its regulatory mechanisms remain incompletely understood.

AMD is a multifactorial disease influenced by both genetic and environmental factors. Genetics association studies have identified a synonymous SNP (rs17810398) within the *DAPL1* gene as associated with AMD [25]. Our previous work demonstrated that Death-associated protein like 1 (DAPL1) is highly expressed in mature RPE cells, where it acts as an inhibitor of proliferation [26]. Furthermore, our recent findings revealed that DAPL1 directly binds the transcription factor E2F4, leading to the upregulation of MITF and its downstream targets NRF2 and PGC1α. This establishes DAPL1 as a regulator of the RPE antioxidant defense system [27]. These collective findings suggest DAPL1 plays a critical role in regulating antioxidant defenses during age-related retinal degeneration. However, the age-related pathologies in *Dapl1-/-* mice remain incompletely characterized. For instance, in our previous study, we observed punctate light spots in aged *Dapl1-/-* mice, resembling AMD features, but the underlying mechanism was unknown.

In this study, we found that autophagy was impaired in *Dapl1-/-* RPE, characterized by decreased levels of ATG16 and Beclin1, alongside increased level of mTOR. Mechanistically, we demonstrate that DAPL1 suppresses the expression of E2F1 (E2F transcription factor 1) and c-MYC (MYC proto-oncogene, bHLH transcription factor). This suppression subsequently inhibits ATG16 and Beclin1 expression and increases mTOR expression, ultimately regulating RPE cell autophagy through DAPK1 (Death associated protein kinase 1). Taken together, our findings demonstrate that DAPL1 acts as a novel regulator of autophagy in RPE cells.

Deficiency in *Dapl1* in mice increases susceptibility to age-dependent retinal pathologies./l

## Materials and methods

### Animal studies

Protocols used for all animal studies were approved by the Wenzhou Medical University Animal Care and Use Committee and performed in compliance with the Association for Research in Vision and Ophthalmology (ARVO) Statement on the use of Animals in Ophthalmic and Vision Research (WYDW2019-0353). C57BL/6J mice were obtained from The Jackson Laboratory and were maintained in the specific pathogen-free facility of the Wenzhou Medical University, China. CRISPR/Cas9 mediated *Dapl1* knockout mice were generated as previously described [26,27]. All mice were given a standard diet and had free access to food and water at all times.

### Cell culture

The ARPE-19 cell line was purchased from ATCC and cultured in DMEM/F12 medium (*Gibco, Eggenstein, Germany*) supplemented with 10% FBS, 100 U/ml of penicillin, and 100 mg/ml of streptomycin. DAPL1-overexpressing ARPE-19 cells (Lv-DAPL1) and ARPE19 (Lv-EGFP) cells were produced as described in the previous work [26,27].

### Reagents and antibodies

Primary antibodies against mTOR, ATG5, ATG16, Beclin1, E2F1, c-MYC, and GAPDH were obtained from Cell Signaling technology (*Beverly, MA*), DAPL1, HA and LC3I/II were from Abcam (*Cambridge, MA*), DAPK1 and GFAP were from Sigma Aldrich (St. Louis, MO), ZO-1 was from Invitrogen (Waltham, MA), Rhodopsin and Opsin were from Millipore (Billerica, MA), IBA1 was from WAKO (Richmood, VA). The goat anti-mouse IgG alexa Fluor® 488 (Life Technologies, A11029) and goat anti-rabbit IgG alexa Fluor® 594 (Life Technologies, A11037) were purchased from Life Technologies.

### Real-Time Quantitative PCR, Western blot analysis

The methods for real-time PCR and western blotting were as described previously [26.27]. Primers for genes, including *DAPK1*, *DAPL1*, *E2F1* and *c-MYC* were synthesized in by Gene Pharma (Shanghai, China). The primer sequences are listed in Supplementary file (Table S1).

### siRNA knock down

ARPE-19 cells were plated in 6-well culture plates at a density of 2×10^5 cells/well. Cells were maintained in a humidified 37°C incubator with 5% CO₂ until reaching 70% confluency (∼24 h). For siRNA-mediated gene silencing, two independent siRNAs targeting human *E2F1*, *c-MYC* transcripts, along with scrambled non-targeting control siRNA, were designed and validated by Gene Pharma Co (sequences in Supplementary Table S2). Transfection complexes were prepared by separately diluting 5 μL of siRNA and 5 μL Lipofectamine™ 2000 in 95 μL serum-free high glucose DMEM medium, followed by 5 min equilibration. The lipid-siRNA complexes were incubated for 20 min at room temperature, and subsequently administered dropwise to cells in serum-free high glucose DMEM medium(final siRNA concentration: 100 nM). Following 6 h transfection under standard culture conditions, the transfection medium was aspirated and replaced with complete high glucose medium containing 10% FBS. Post-transfection incubation proceeded for 24 h to allow target protein knockdown and the following experiment such as western blot was performed.

### Lentivirus infection

DAPL1-HA was constructed into the vector of TG005, which was purchase from the Bi’ang Biomedical Technology Co., Ltd (Shanghai, China). DAPL1-HA cDNA was amplified using the primers of DAPL1-HA-F: ATACGCGTATGGCGTATCCCTACGACGTGCCCGATTATGCCGCAAATGAAGTGCAAGA; DAPL1-HA-R: CGACTAGTTTAGGCATAATCGGGCACGTCGTAGGGATACGCACATTTTCGAGGCTGCT. DAPL1-Del-HA cDNA was amplified using the primers of DAPL1-Del-HA-F: AAGCTGGAATGCAAGAAATTG; DAPL1-Del-HA-R: CAATTTCTTGCATTCCTCCAGCTT. Lentiviral particles for DAPK1 (LPP-H0465-Lv242-100), E2F1, E2F4 and c-MYC (LPP-Z2845-Lv105-100) were purchased from Gene Copoeia Company. For infection, viruses were suspended in serum-free medium and added to REFs at a multiplicity of infection (MOI) of 1,000. After 30 min incubation, fresh medium was added to the cells. After culturing for 24 h, the following experiment such as western blot was performed. For viral infection, lyophilized viral particles were reconstituted in serum-free DMEM and diluted to achieve the desired multiplicity of infection (MOI=1,000). Confluent REF cultures were inoculated with the viral suspension, followed by a 30-min adsorption period at 37°C in a 5% CO2 humidified incubator. Post-infection incubation proceeded for 24 h under standard culture condition. Protein lysates were prepared using RIPA buffer and clarified supernatants were subjected to western blot analysis as previously described [27].

### Fundus photography and optical coherence tomography (OCT)

Mice were anaesthetized with ketamine (90 mg/kg) and xylazine (8 mg/kg). The iris was dilated with 1% Mydriacil and the corneas kept moist with saline solution during photography. The mouse was positioned on a heating pad during the procedure to maintain the body temperature. Photographs were made with a Kowa small animal fundus camera (2.5 megapixels) equipped with a 66 diopter supplemental lens (Kowa American Corporation, Torrance, CA).

### Tissue collection and retinal histological analysis

Following humane euthanasia via anesthetic overdose, bilateral ocular globes were immediately enucleated for multimodal analysis. Tissue processing protocols were stratified according to experimental requirements: (1) For histopathological evaluation, eyes were fixed in 4% paraformaldehyde (PFA) for 24 h at 4°C prior to paraffin embedding and microtome sectioning (5 μm). (2) Eye samples were embedded in OCT (Optical Cutting Temperature) compound at - 80°C for immunofluorescence studies. (3) Immunofluorescence protocols included sequential antigen retrieval (0.1% Triton X-100, 30 min) and blocking with 1% bovine serum albumin (BSA) for 1 h at room temperature. RPE flat mounts were probed with anti-ZO-1 primary antibody in humidified chambers at 4°C overnight, followed by secondary antibody incubation for 2 h. Parallel retinal lysates prepared in RIPA buffer containing protease inhibitors were subjected to western blot analysis using standard SDS-PAGE protocols. Lipid accumulation was quantified through Oil Red O staining of cryosections counterstained with hematoxylin.

### Statistical Analysis

Data were presented as the mean ± SEM. The statistical significance of differences between groups was obtained using one-way ANOVA in GraphPad (*San Diego, CA*) . Differences were considered to be significant at P < 0.05.

## Results

### Aged *Dapl1-/-* mice retinas display age-dependent retinal pathologies

In our previous study, we identified age-related functional and structural abnormities in the retinas of aged *Dapl1-/-* mice [27]. Intriguingly, additional pathologies were observed in these aged knockout mice. Fundus photography revealed punctate light deposits in the eyes of 18-month-old *Dapl1-/-* mice (Fig 1A left, arrows). Optical coherence tomography (OCT) confirmed the presence of these deposits located between the outer nuclear layer (ONL) and the RPE (Fig 1A, right, arrows). These deposits likely represent accumulations of autofluorescent cells and lipid aggregates within the aged *Dapl1-/-* retina. Supporting this, abnormal cells staining purple with hematoxylin were observed at the ONL-RPE interface (Fig 1B left). Furthermore, both these cells and the adjacent RPE exhibited autofluorescence (Fig 1B, middle), resembling that of lipofuscin and lipid-rich aggregates known to accumulate in aging tissues and implicated in age-dependent retinal pathologies [1,28]. Oil Red O staining confirmed the lipid-rich nature of these deposits within the retina (Fig 1B, right). To determine the identity of the autofluorescent cells, immunostaining was performed. These subretinal autofluorescent cells were positive for IBA1, a microglia marker, in the 18-month-old *Dapl1-/-* retina (Fig 1C). This finding suggests that stress originating within the RPE layer may trigger microglial migration towards this site [1]. In conclusion, aged *Dapl1-/-* mice develop age-dependent retinal pathologies characterized by the accumulation of lipid-rich, autofluorescent deposits associated with microglial infiltration.

**Fig 1.**
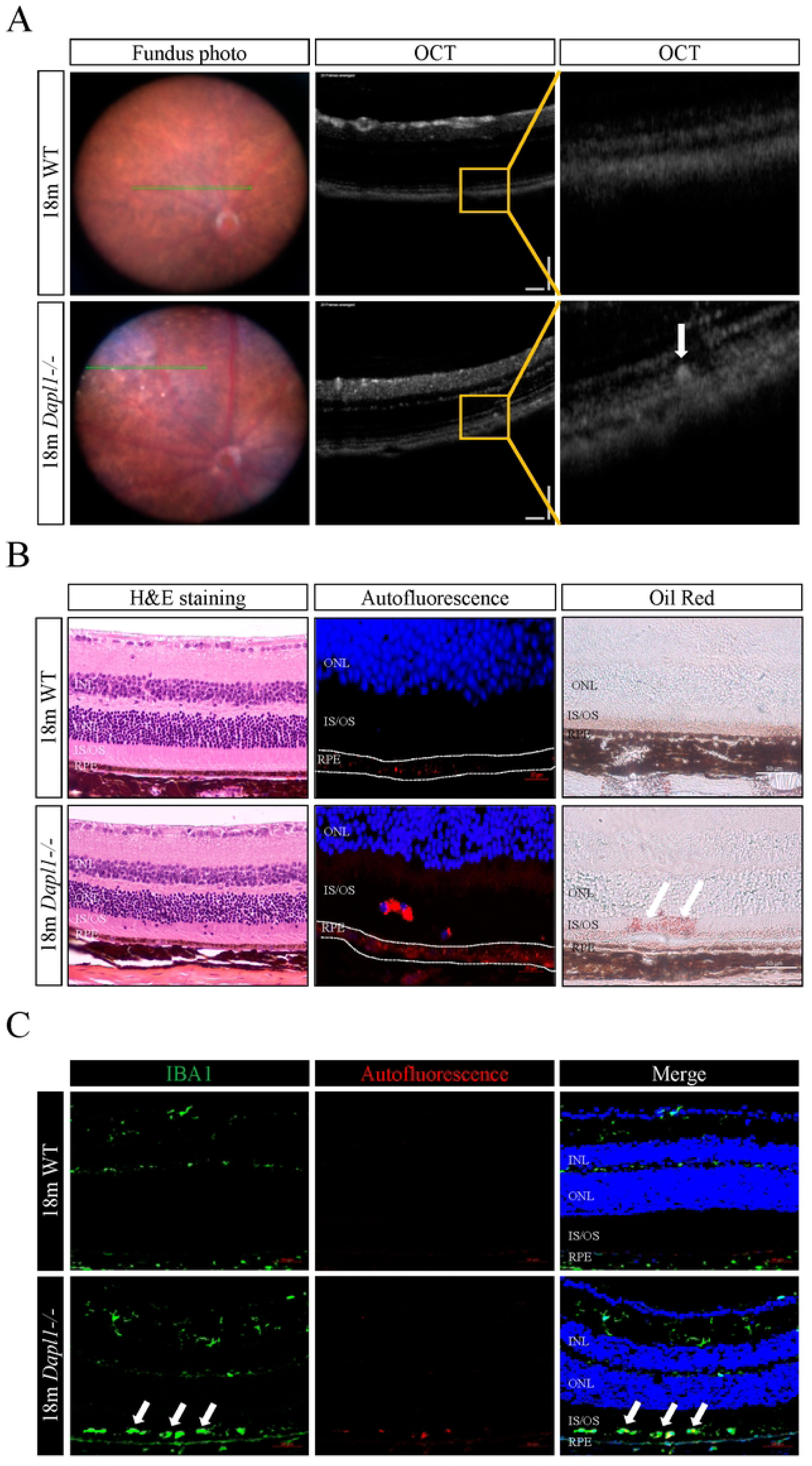
Aged *Dapl1-/-* mice retinas display age-dependent retinal pathologies. (A) Fundus photography of 18-month-old *Dapl1-/-* mice reveals numerous white spots (arrows). OCT analysis confirms the presence of these deposits (arrows). (B) Retinal pathology in 18-month-old *Dapl1-/-* mice. (B, left) H&E staining shows clusters of nuclei within the subretinal space between the ONL and RPE. Scale bar, 10 µm. (B, middle) Autofluorescence is observed in the IS/OS layer and RPE layer. Scale bar, 20 µm. (B, right) Oil Red O staining indicates lipid droplet accumulation (arrows) between the ONL and RPE. Scale bar, 50 µm. (C) IBA1 immunostaining identifies microglia. Positive cells (arrows) suggest microglial migration into the subretinal space (between ONL and RPE). Scale bar, 50 µm, n=6, **P<0.01.

### Aged *Dapl1-/-* mice exhibit retinal and RPE pathology

Retinal and RPE cell damage are key contributors to age-dependent retinal pathologies. As shown in the data, up-regulation of GFAP (glial fibrillary acidic protein) was observed in 18-month-old *Dapl1-/-* mice compared to the wild-type (WT) controls (Fig 2A), indicating significantly increased retinal stress. Consistent with this finding, expression levels of the photoreceptor markers Rhodopsin (rods) and Opsin (cones) were also reduced in aged *Dapl1-/-* mice (Fig 2B). Furthermore, we assessed RPE structure using ZO-1 (tight junction protein 1, TJP1) staining in RPE whole mounts from 18-month-old mice. While WT mice exhibited a regular hexagonal RPE mosaic with intact junctions, *Dapl1-/-* mice showed enlarged, multinucleated RPE cells (open arrowheads) and disrupted tight junctions (solid arrows) (Fig 2C). These results demonstrate that loss of DAPL1 in mice leads to photoreceptor and RPE damage, culminating in age-dependent retinopathies.

**Fig 2.**
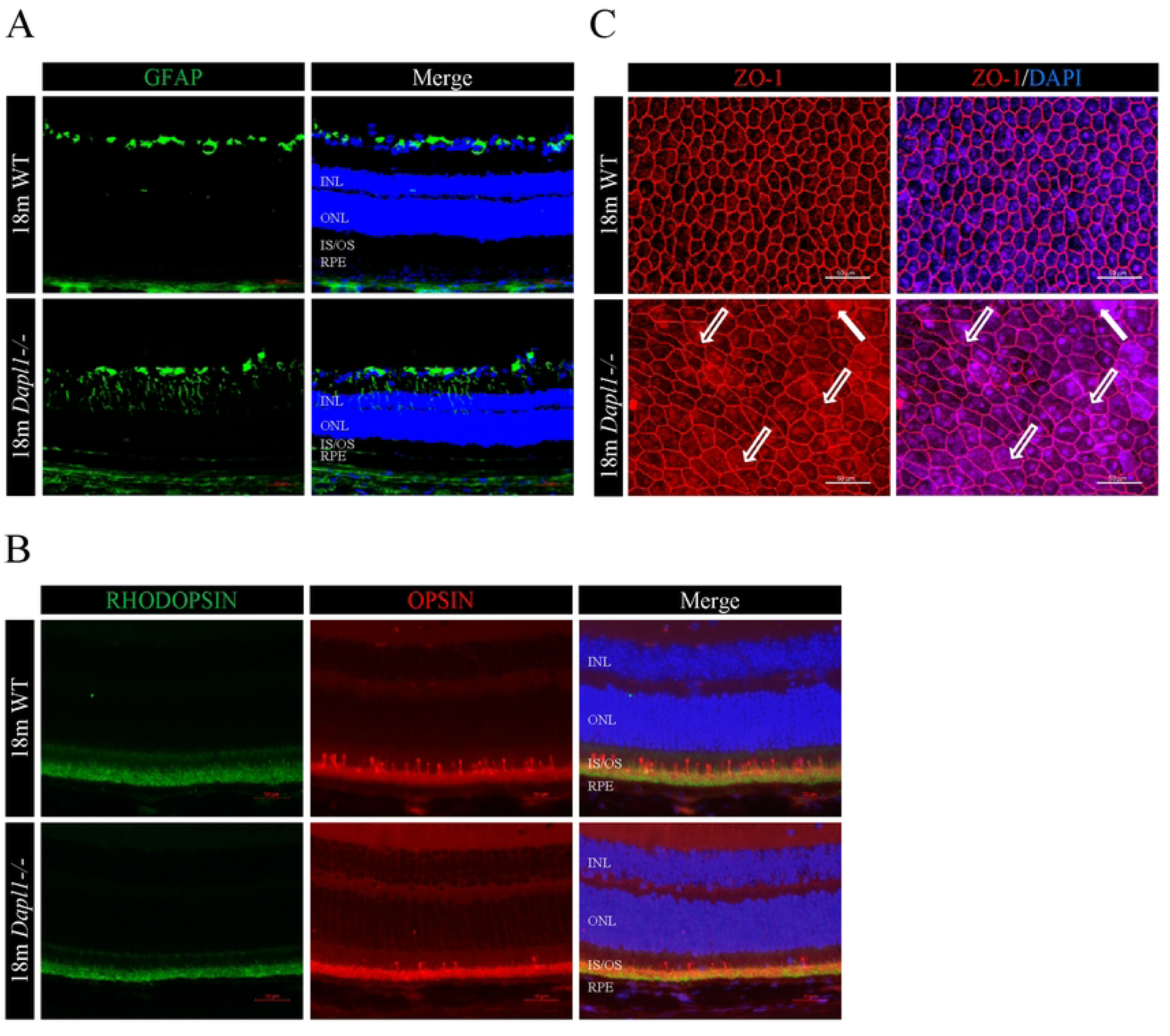
Aged *Dapl1-/-* mice exhibit retinal and RPE pathology. (A) Increased GFAP levels in the retinas of 18-month-old *Dapl1-/-* mice. (B) Reduced expression of photoreceptor markers Rhodopsin and Opsin in the 18-month-old *Dapl1-/-* retinas. (C) ZO-1 immunostaining of RPE flat-mounts. While WT mice show a regular hexagonal RPE mosaic, 18-month-old *Dapl1-/-* mice exhibit multinucleated RPE cells (hollow arrowheads) and disrupted tight junctions (solid arrows). Scale bar, 50µm, n=6.

### DAPL1 regulates autophagy in RPE cells

The above results demonstrated that loss of DAPL1 in the RPE increases susceptibility to age-dependent retinal pathologies in mice. However, our data did not identify the specific RPE cellular function regulated by DAPL1. Given that RPE autophagy deficiency is linked to age-related retinal degeneration [17–18], we investigated whether autophagy impairment underlies the pathologies in *Dapl1-/-* mice. LC3 immunostaining of RPE/retina sections from 18-month-old mice revealed significantly weaker LC3 signal in *Dapl1-/-* retinas compared to WT (Fig 3A), suggesting impaired autophagy. Consistent with this, protein levels of autophagy-related markers ATG16 and Beclin1 were decreased in *Dapl1-/-* RPE, while the negative regulator mTOR was increased (Fig 3B, C). These findings indicate that DAPL1 deficiency suppresses autophagy in RPE cells. To further validate DAPL1’s role, we overexpressed DAPL1 in ARPE-19 cells (Lv-DAPL1) [26]. DAPL1 overexpression increased ATG16 and Beclin1 protein levels while reducing mTOR (Fig 3D-G). Western blot analysis showed elevated LC3-II levels in Lv-DAPL1 cells versus Lv-EGFP controls under both normal and nutrient-starved conditions (Fig 3H, I). To assess autophagic flux, we treated cells with NH_4_Cl to inhibit autophagosome degradation. LC3 immunostaining revealed significantly more LC3-positive puncta in DAPL1-overexpressing cells (Fig 3J, K). Collectively, these results demonstrate that DAPL1 promotes autophagy in RPE cells.

**Fig 3.**
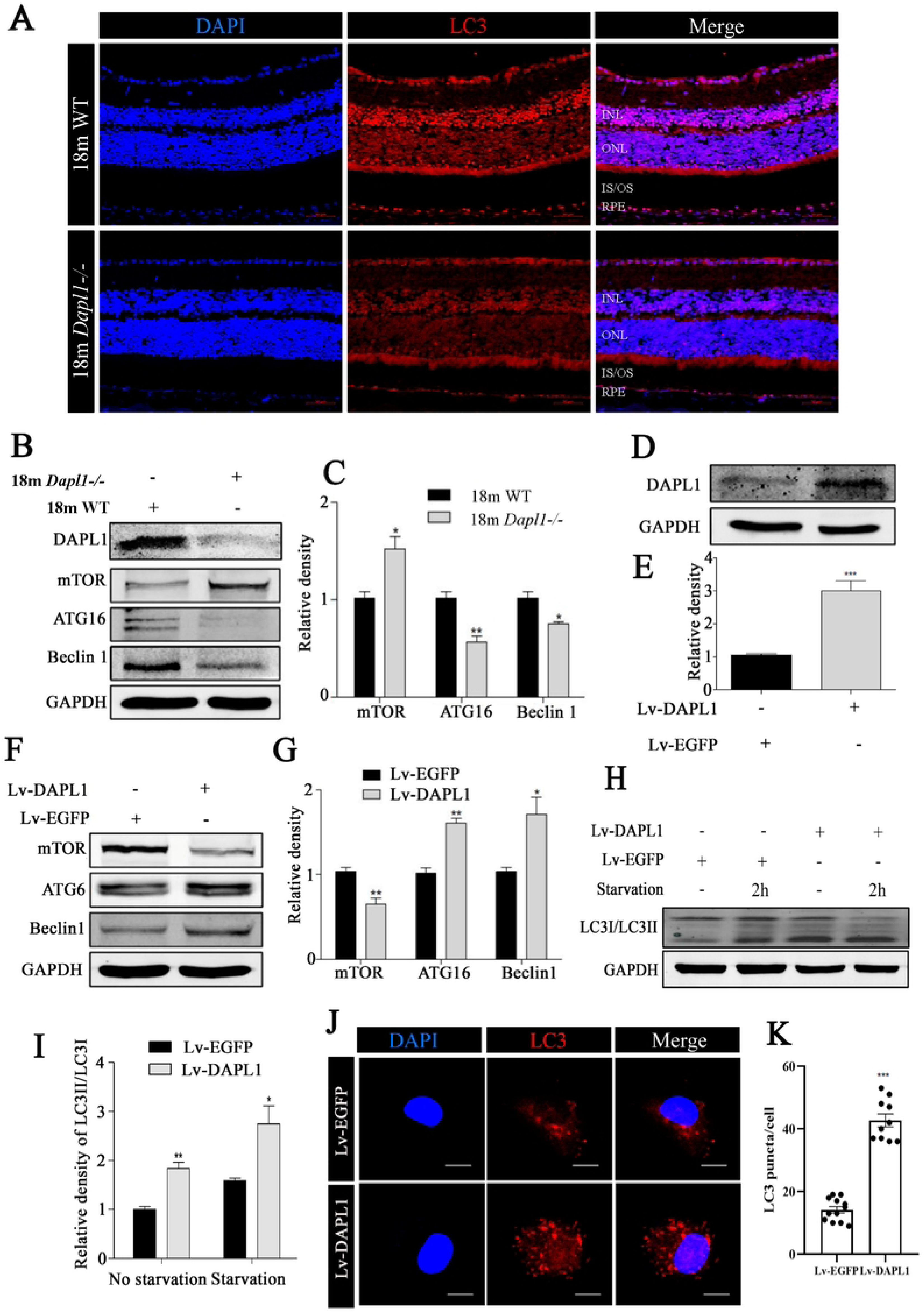
DAPL1 regulates autophagy in RPE cells. (A) Representative LC3 immunostaining in RPE-retina sections from18-month-old *Dapl1-/-* and WT mice. Scale bar, 50µm, n=6. (B) Western blot analysis of mTOR, ATG16, and Beclin1 protein levels in RPE cells from 18-month-old mice. (C) Quantification of western blot bands shown in (B). (D) Western blot confirming DAPL1 overexpression in ARPE-19 cells transduced with Lv-DAPL1. (E) Quantification of DAPL1 protein levels from (D). (F) Western blot analysis of mTOR, ATG16, and Beclin1 protein levels in ARPE-19 cells transduced with Lv-EGFP or Lv-DAPL1. (G) Quantification of western blot bands shown in (F). (H) Western blot analysis of LC3-I/II conversion in ARPE-19 cells under normal and nutrient-starved conditions. (I) Quantification of LC3-II protein levels from (H), n=3. (J) LC3 immunostaining in ARPE-19 cells transduced with Lv-EGFP or Lv-DAPL1 following treatment with NH₄Cl under nutrient starvation to assess autophagic flux. Scale bar, 10µm, n=10. (K) Quantification of LC3-positive puncta per cell from ***P<0.001, **P<0.01, *P<0.05.

### DAPL1 regulates RPE cell autophagy partially through DAPK1

The above results demonstrated that DAPL1 regulates autophagy in RPE cells, but the underlying mechanisms remain unclear. Our previous work established that DAPL1 decreases E2F1 protein levels in ARPE-19 cells [26], which could inhibit autophagy [29]. Furthermore, DAPL1 binds E2F4 to suppress c-MYC expression [27]. Notably, both E2F1 and c-MYC regulate DAPK1 (Death-associated protein kinase 1) expression [30], and DAPK1 plays a critical role in autophagy regulation [31]. To determine whether DAPL1 regulates autophagy via the E2F1/c-MYC-DAPK1 axis, we first analyzed DAPK1 expression in *Dapl1-/-* mouse RPE and DAPL1-overexpressing ARPE-19 cells. DAPK1 mRNA and protein levels were significantly elevated in *Dapl1-/-* RPE (Fig 4A, 4C-D). Conversely, DAPK1 was reduced in DAPL1-overexpressing cells (Fig 4B, 4E-F). Overexpression of DAPK1 in ARPE-19 cells increased mTOR while decreasing Beclin1 and ATG16 levels (Fig 4G-H), indicating DAPK1 suppresses autophagy. We next tested whether DAPK1 overexpression could rescue DAPL1-induced autophagy. Co-overexpression of DAPK1 partially reversed the effects of DAPL1 on key autophagy markers: it attenuated DAPL1-mediated increases in LC3-II/LC3-I ratio, ATG16, and Beclin1, while counteracting DAPL1-induced mTOR suppression (Fig 4I-L). These findings demonstrate that DAPL1 promotes RPE autophagy partially by downregulating DAPK1 expression via the E2F1/c-MYC axis.

**Fig 4.**
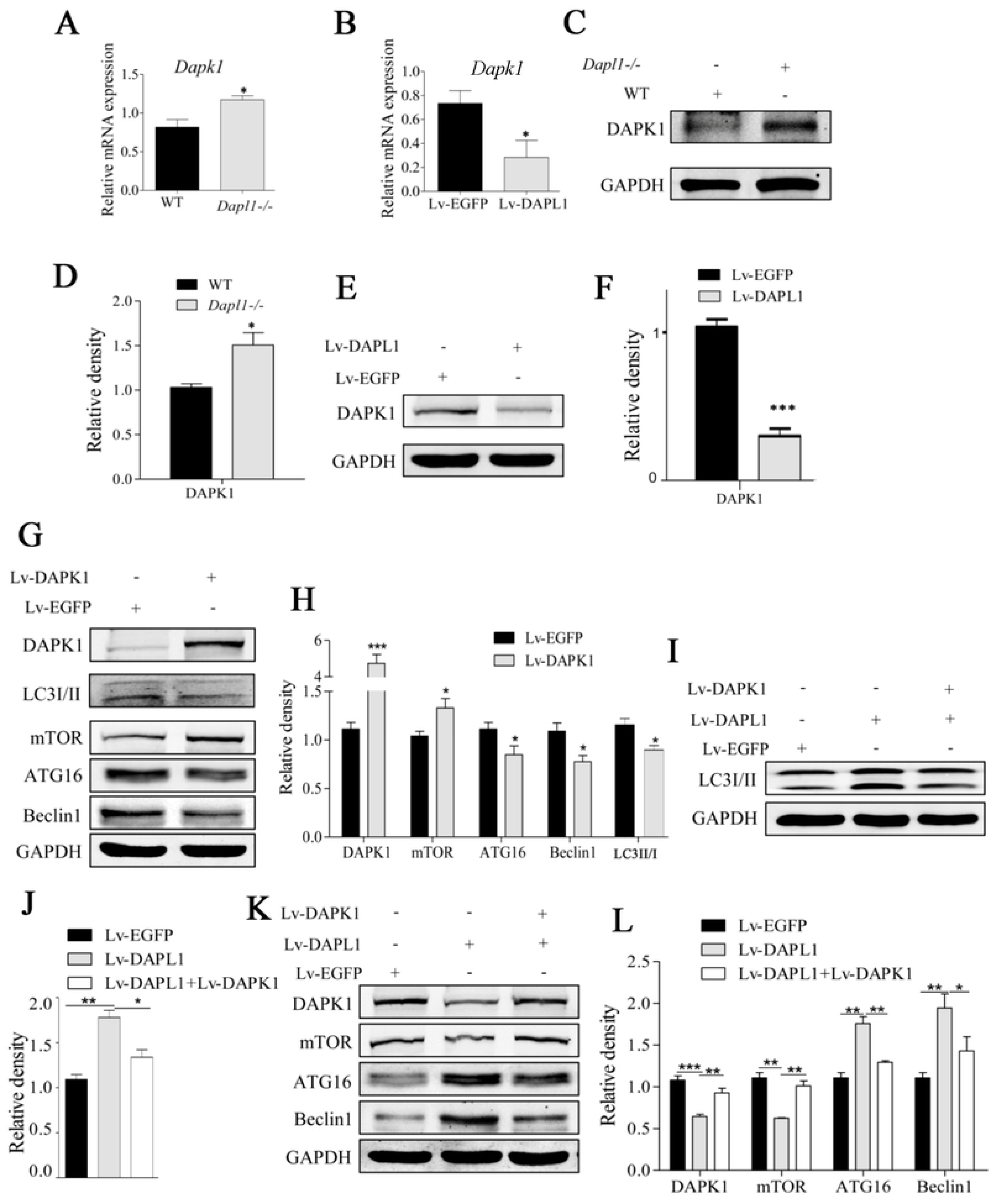
DAPL1 regulates RPE cell autophagy partially through DAPK1. (A) *Dapk1* mRNA levels in *Dapl1-/-* and WT RPE cells (qRT-PCR). (B) *Dapk1* mRNA levels in ARPE-19 cells transduced with Lv-EGFP or Lv-DAPL1 (qRT-PCR). (C) Representative Western blot of DAPK1 protein in *Dapl1-/-* and WT RPE. (D) Quantification of DAPK1 protein levels from (C). (E) Representative Western blot of DAPK1 protein in ARPE-19 cells transduced with Lv-EGFP or Lv-DAPL1. (F) Quantification of DAPK1 protein levels from (E). (G) Representative Western blot of mTOR, LC3-II, ATG16, and Beclin1 in Lv-DAPK1-transduced and Lv-EGFP-transduced ARPE-19 cells. (H) Quantification of protein levels from (G). (I) Representative Western blot of LC3-I/II in Lv-DAPL1 cells with/without DAPK1 co-overexpression. (J) Quantification of LC3-II/LC3-I ratio from (I). (K) Representative Western blot of mTOR, ATG16, and Beclin1 in Lv-DAPL1 cells with/without DAPK1 co-overexpression. (L) Quantification of protein levels from (J) n=3, **P<0.01, *P<0.05.

### DAPL1 regulates DAPK1 expression through E2F1 and c-MYC

The above data demonstrated that DAPL1 regulates autophagy by inhibiting DAPK1 expression in ARPE-19 cells. However, it remained unclear whether DAPL1 suppresses DAPK1 via E2F1/c-MYC. To address this, we analyzed E2F1 and c-MYC protein levels. Both E2F1 and c-MYC were elevated in *Dapl1-/-* RPE cells but reduced in DAPL1-overexpressing ARPE-19 cells (Fig 5A, B). Furthermore, knocking down E2F1 or c-MYC in ARPE-19 cells could decrease the protein level of DAPK1 (Fig 5C, D), while overexpressing E2F1 or c-MYC increased DAPK1 levels (Fig 5E, F). Finally, we tested whether E2F1/c-MYC overexpression could counteract DAPL1-mediated effects in ARPE-19+Lv-DAPL1 cells. Western blotting showed that overexpressing E2F1 or c-MYC in ARPE-19+Lv-DAPL1 cells increased DAPK1 expression, and reversed DAPL1-induced changes in mTOR, Beclin1 and ATG16 (Fig 5G, H). These results demonstrate that DAPL1 regulates DAPK1 expression through the E2F1/c-MYC pathway.

**Fig 5.**
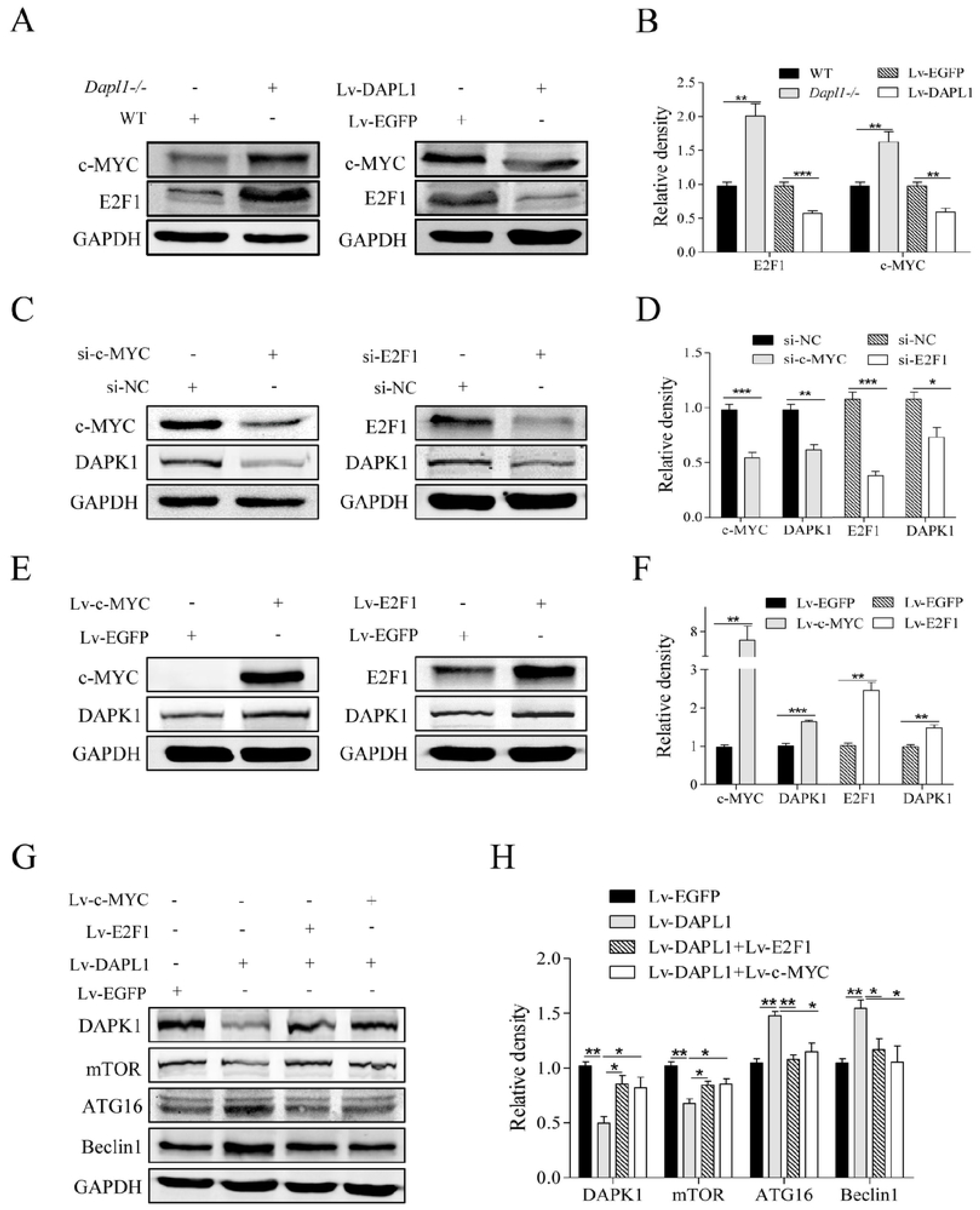
DAPL1 regulates DAPK1 expression through E2F1 and c-MYC. (A) Representative Western blots of E2F1 and c-MYC in *Dapl1-/-* RPE and DAPL1-overexpressing ARPE-19 cells. (B) Quantification of protein levels from (A). (C) Representative Western blots showing DAPK1 reduction in ARPE-19 cells following E2F1 or c-MYC knockdown. (D) Quantification of DAPK1 levels from (C). (E) Representative Western blots showing DAPK1 elevation in ARPE-19 cells overexpressing E2F1 or c-MYC. (F) Quantification of DAPK1 levels from (E). (G) Representative Western blots demonstrating that E2F1/c-MYC overexpression in DAPL1-overexpressing ARPE-19 cells reverses DAPK1 levels and rescues DAPL1-mediated changes in mTOR, ATG16, and Beclin1. (H) Quantification of protein levels from (G). n=3, ***P<0.001, **P<0.01, *P<0.05.

**Fig 6.**
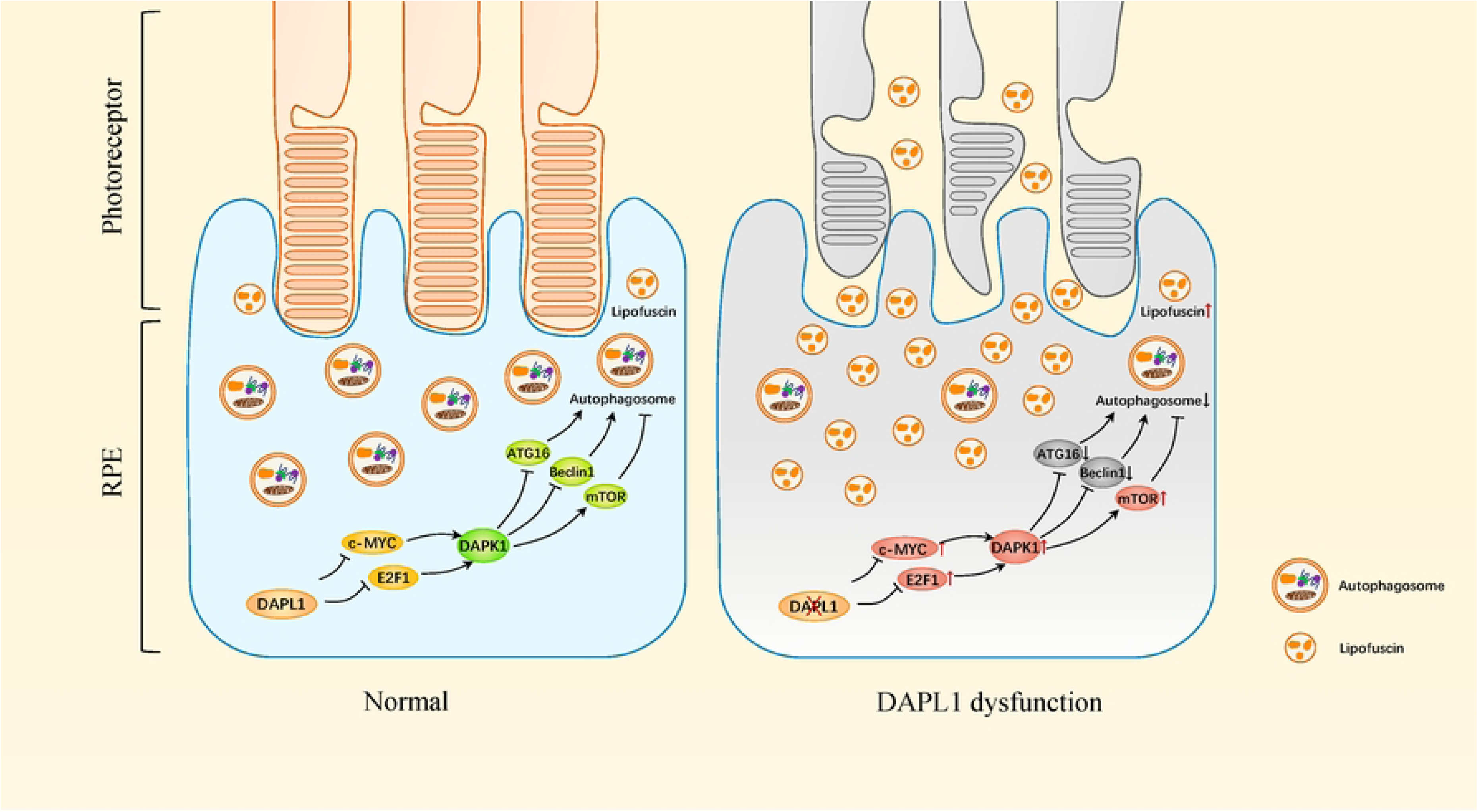
Model: DAPL1 regulates RPE autophagy through the E2F1/c-MYC-DAPK1 axis. DAPL1 maintains retinal homeostasis by promoting autophagy through suppression of the E2F1/c-MYC-DAPK1 pathway. This regulation modulates key autophagy components (ATG16/Beclin1/mTOR). Loss of DAPL1 in RPE cells inhibits autophagy, contributing to age-dependent retinal pathology.

### Graphical summary

## Discussion

Age-related retinopathies, particularly age-related macular degeneration (AMD), represent the leading cause of irreversible vision loss in geriatric populations worldwide. Autophagic dysfunction has been suggested in a broad range of neurodegenerative diseases, including Alzheimer, Huntington diseases and AMD [32,33], but the precise molecular mechanisms governing autophagy regulation in RPE cells remain incompletely characterized. Our findings reveal a novel regulatory axis wherein DAPL1 coordinates autophagic flux through the E2F1/c-MYC-DAPK1-mTOR/ATG16/Beclin1 signaling cascade in RPE cells. Crucially, *Dapl1*-deficient murine models exhibit impairment of RPE autophagic capacity and age-dependent retinal pathologies.

Aging and RPE dysfunction are two main causing factors in age-related retinal degeneration. Oxidized proteins, lipids and damaged mitochondria are increased in the aging tissues [34,35], together with the environmental risk factors, such as smoking and excessive light exposure, which contribute to the development of retinal degeneration. It might be one of the reasons why *Dapl1-/-* mice have normal retinal structure and visual function in the young age, but display age-dependent retinal pathologies in the old age.

RPE cells constitute the cornerstone of retinal homeostasis through two primary mechanisms: (1) antioxidant defense via glutathione redox cycling and Nrf2-mediated pathways, and (2) circadian-regulated phagocytosis of shed photoreceptor outer segments. Retinal oxidative damage has been identified as one of the major causative factors of AMD. Deficiency in RPE cells antioxidant ability is critically implicated in the pathogenesis of AMD, while enhancement of the antioxidant capability of the RPE cells could protect the neural retina from oxidative damage [4,36,37]. Upregulation of autophagy in RPE cells reduces oxidative stress-induced generation of reactive oxygen species (ROS), whereas autophagy inhibition increases ROS generation, promotes lipofuscin accumulation, and heightens susceptibility to oxidative stress-induced damage [18]. Building on this, our recent work demonstrates that DAPL1 serves as an antioxidant regulator in RPE cells, modulating the MYC-MITF-NRF2/PGC1α axis [27]. Our current findings identify that *Dapl1* deficiency in mice inhibits RPE cell autophagic flux, revealing a novel role for DAPL1 in regulating the RPE autophagy system.

Beyond autophagy, heterophagy—specifically the phagocytosis of shed POS—is highly active in RPE cells. This process occurs in a diurnal rhythm and is essential for photoreceptor renewal and RPE cell survival [38]. Defect of PRE phagocytosis, such as those caused by *MERTK* (MER proto-oncogene, tyrosine kinase) mutations, lead to accumulation of unphagocytosed POS debris, retinal degeneration, and blindness in both humans and animal models [39,40]. Notably, RPE cells utilize autophagy proteins (e.g., LC3, Beclin1) for phagosome formation during LC3-associated phagocytosis (LAP). Consistently, *Atg5* deficiency in RPE cells impairs their capacity to degrade POS, resulting in retinal dysfunction [33,41].

While our findings establish DAPL1 as a critical modulator of autophagic flux in RPE cells, a critical knowledge gap persists regarding its potential involvement in phagocytic regulation.

Our prior investigations revealed that DAPL1 expression is transcriptionally controlled by MITF (Microphthalmia-associated transcription factor) specific isoform MITF-6 in RPE cells [42]. Consistently, *Mitf* deficiency mice exhibit RPE dysfunction and retinal degeneration [8]. While MITF has been shown to regulate autophagy via miR-211 in Hela, HEK293T and melanoma cell lines [43], its role in RPE cell autophagy remains undefined. Here, we demonstrate that DAPL1—a downstream MITF target—positively regulates RPE cell autophagy by promoting ATG16 and Beclin1 expression while inhibiting mTOR. Notably, the mTOR pathway is associated with RPE differentiation and retinal degeneration [44] and can regulate MITF expression through β-catenin [45]. Collectively, our findings suggest a potential positive-feedback loop between DAPL1 and MITF in RPE cells, regulating autophagy and retinal homeostasis.

Cellular proliferation imposes significant demands for energy and reducing power, while proliferation-inducing agents elevate cellular oxidative stress [46]. Autophagy plays critical roles in maintaining cellular homeostasis and alleviating metabolic stress. Notably, autophagy is inhibited during mitosis to safeguard nuclear integrity and genomic stability [47,48]. Our previous work established DAPL1 as a cell proliferation inhibitor in RPE cells [26]. Furthermore, key DAPL1 downstream targets, E2F1 and c-MYC, regulate both autophagy and cell cycle progression [49]. The intricate relationship between DAPL1-mediated regulation of cell proliferation and autophagy in RPE cells remains poorly understood. Building on our current findings, we propose that DAPL1 critically regulates the RPE cell cycle, cellular homeostasis, and metabolic balance—thereby establishing a protective metabolic window for essential quality control processes—which ultimately contributes to age-dependent retinal pathologies.

DAPK1, a calcium/calmodulin (Ca²⁺/CaM)-regulated serine/threonine kinase, is extensively studied in tumor cell death [50], yet its functions in RPE cells remain unexplored. Notably, E2F1 and c-MYC inhibit autophagy in other cell types [29,51]—consistent with our findings—though their role in RPE cell autophagy is undefined. Our mechanistic studies identify a novel transcriptional axis in which DAPK1 expression is upregulated by E2F1 and c-MYC, aligning with observations in embryonic fibroblasts [30]. However, while DAPK1 is widely reported to promote autophagy [52], our data reveal it mediates E2F1/c-MYC-driven autophagy inhibition in RPE cells. This functional divergence suggests DAPK1 may exert context-dependent biological roles.

In summary, this investigation elucidates the mechanistic role of DAPL1—a recognized AMD susceptibility locus—in orchestrating retinal homeostasis through autophagic regulation. Our data establish that DAPL1 serves as a critical nodal regulator of the E2F1/c-MYC→DAPK1→mTOR/ATG16/Beclin1 multi-tiered signaling cascade in RPE cells. Our findings not only enhance the understanding of the regulatory mechanisms of autophagy, but also provides insight into the cellular and molecular mechanisms of DAPL1 deficiency induced age-dependent retinal pathologies. These findings also warrant additional studies to elucidate the functional roles of DAPL1 in RPE function and retinopathies, which might point to the prevention or treatment of age-related retinal diseases.

## Acknowledgements

This work was supported by the National Natural Science Foundation of China (81770946).

## Authors’ contributions

### Conceived and designed the experiments

Ling Hou, Huaicheng Chen.

### Performed the experiments

Huaicheng Chen, Qiufan Tan.

### Data analyzation

Huaicheng Chen, Qiufan Tan, Yishan Hu, Liping Liu and Xiaoxin Liang.

### Contributed reagents/materials/analysis tools

### Wrote the manuscript

Huaicheng Chen, Qiufan Tan

### Revised the manuscript

Huaicheng Chen, Ling Hou

### Data availability

Data will be made available on request.

## Supplementary data

**Table S1:**
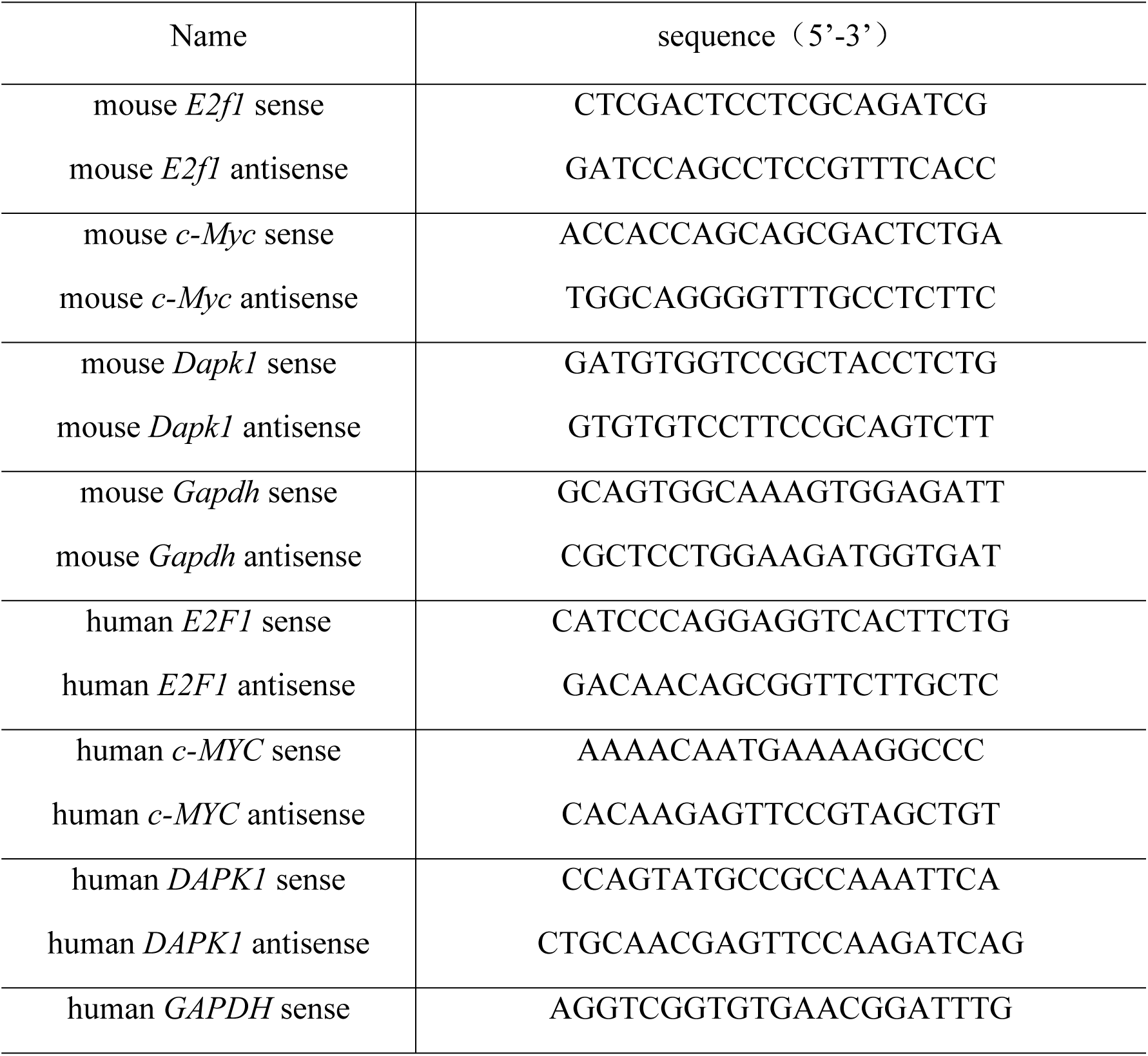
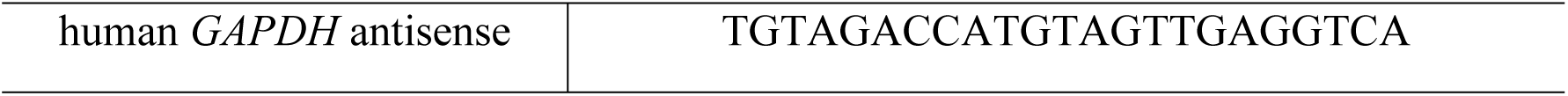
Sequence of primers.

**Table S2:**
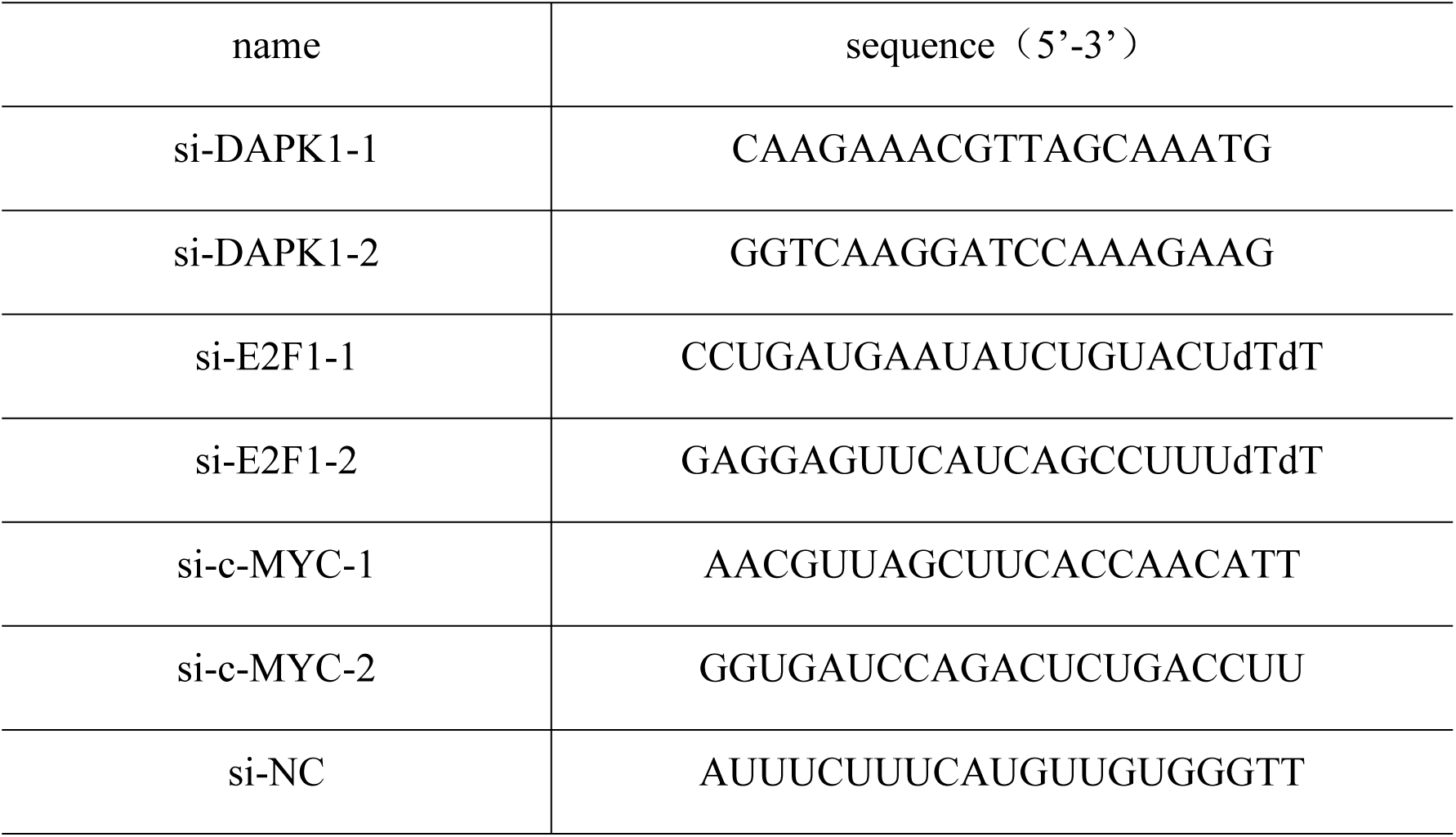
Sequences of siRNAs:

## Abbreviations

AMD: age-related macular degeneration
ANOVA: analysis of variance
ATG16: autophagy related 16
c-MYC: MYC proto-oncogene, bHLH transcription factor
DAPL1: Death associated protein like 1
DAPK1: Death associated protein kinase 1
E2F1: E2F transcription factor 1
E2F4: E2F transcription factor 4
GFAP: glial fibrillary acidic protein
INL: inner nuclear layer
IS/OS: inner segment/outer segment
mTOR: mechanistic target of rapamycin
OCT: optical coherence tomography
ONL: outer nuclear layer
PKC: protein kinase C
ROS: reactive oxygen species
RPE: Retinal pigment epithelial cell
WT: wild type.

